# Ultrasound-enhanced retinal delivery of engineered viral vectors

**DOI:** 10.64898/2026.01.04.697446

**Authors:** Schuyler Link, Kara Fan, Jesus Adame, Caroline Keehn, Benjamin J. Frankfort, Jerzy O. Szablowski

## Abstract

The goal of this study was to examine Focused Ultrasound Blood-Retina Barrier Opening (FUS-BRBO) as a method to deliver gene therapy to the eye. We compared AAV.FUS.3, an engineered variant optimized for FUS-enhanced delivery to the central nervous system, and AAV9 for their utility as retinal gene vectors. To achieve this, mouse eyes were insonated by 4.667 MHz FUS in a custom-designed nosecone and vectors administered intravenously. We found 2.1-fold improvement of transduction of AAV.FUS.3 over AAV9 in the insonated area and 19.5-fold improvement over the uninsonated retina. Within that area, the ganglion cell layer showed the highest levels of transduction at 22% for AAV.FUS.3. We observed no gross tissue damage, change in retinal layer thickness, or increase in inflammation markers within the retina. FUS-BRBO has the potential to non-surgically deliver AAV-based vectors to spatially-defined regions of the retina. AAV.FUS.3 showed significantly improved transduction over AAV9 at the same dose. Further research to improve efficiency of delivery to the outer retina could expand the utility of FUS-BRBO.

## INTRODUCTION

Delivering drugs to the retina is difficult due to several barriers within and around the eye: topical drugs are prevented from reaching the posterior eye by the corneal epithelium^1–3^, intravitreal drugs may be blocked at the inner limiting membrane between the vitreous and retina^4,5^ and the blood-retina barrier (BRB) prevents therapeutics from entering the retina via systemic circulation^1,6,7^. In this latter case, the vasculature of the retina is reinforced with tight junctions, constituting the inner BRB^6–8^. To circumvent the BRB, direct injection into the vitreous is often used for large molecule therapeutics like biologics^9^. While effective, this invasive procedure has potential serious adverse outcomes including endophthalmitis, retinal detachment, and cataracts^1,10,11^. Furthermore, intravitreal injections can be uncomfortable and even painful, which diminishes patient adherence^12^. Thus, opportunities exist to improve drug delivery such that both procedural risk and discomfort are simultaneously reduced.

Focused Ultrasound (**FUS**) Blood-Brain Barrier opening (**FUS-BBBO**) is a novel drug delivery method to the central nervous system (**CNS**) that has been investigated over the last two decades^13–16^. When applied to systemically administered ultrasound contrast, FUS causes oscillation of microbubble size which exerts mechanical pressure and disrupts the tight junctions that line the vasculature of the brain^17–19^. This allows larger macromolecules such as biologics and gene therapies to access targeted regions^20,21^. This procedure has been demonstrated as safe in a range of animal^22–28^ and clinical studies^22,29–33^, allowing the passage of genetic vectors with therapeutic payloads^21,22^.

As the BBB and BRB are similar, it is not surprising that two groups have demonstrated focused ultrasound blood-retinal barrier opening (**FUS-BRBO**). Park et al. found MRI contrast signal increase in the retina following insonation which could no longer be detected three hours after BRBO^34^. However, histology showed damage to the retina^34^ similar to that which is seen in FUS-associated damage in the brain at high pressure^35,36^. Touahri et al. refined the BRBO procedure and demonstrated the possibility of therapeutic delivery through a systemically administered AAV crossing the BRB^37^. The authors used AAV2/8 to transduce Muller glia and astrocytes in the inner retina, but did not investigate neuronal transduction^37^. Both of these studies used rats, MRI guidance, and ultrasound frequencies of 0.69^34^ and 1.1^37^ MHz, similar to what is used for FUS in rodent brains^34,37^.

In this study, we improved techniques for FUS-BRBO to develop a novel method for gene therapy delivery to the eye. Using high frequency (4.667 MHz) focused ultrasound, we compared two AAV serotypes: an engineered variant optimized for FUS-enhanced delivery to the central nervous system, AAV.FUS.3, and the naturally occurring AAV9, for their utility as retinal gene vectors.

## RESULTS

### Design Of The 3D-Printed Holder for Insonation of The Eye

We adapted a stereotactic-guided FUS (RK50, FUS Instruments) to insonate the eye. We designed a nosecone which rotates the mouse on its side by 70° and pitches the mouse forward by 15° (**Fig. 1A**). This nosecone was accompanied by caps for stereotactic ear bars to support the head and neck (**Fig 1B**). The nosecone, baseplate, and ear-bar covers were 3D printed in polylactic acid (**Fig. 1C**). The orientation of the mouse in the nosecone places the nasal retina 2.8mm inferior to the center of the cornea, when calculating the eye size with an axial length of 3.3 mm^38^ (**Fig. 1D**). The 3-dimensional designs are available for download as Supplementary Files 1-4.

**Figure 1.**
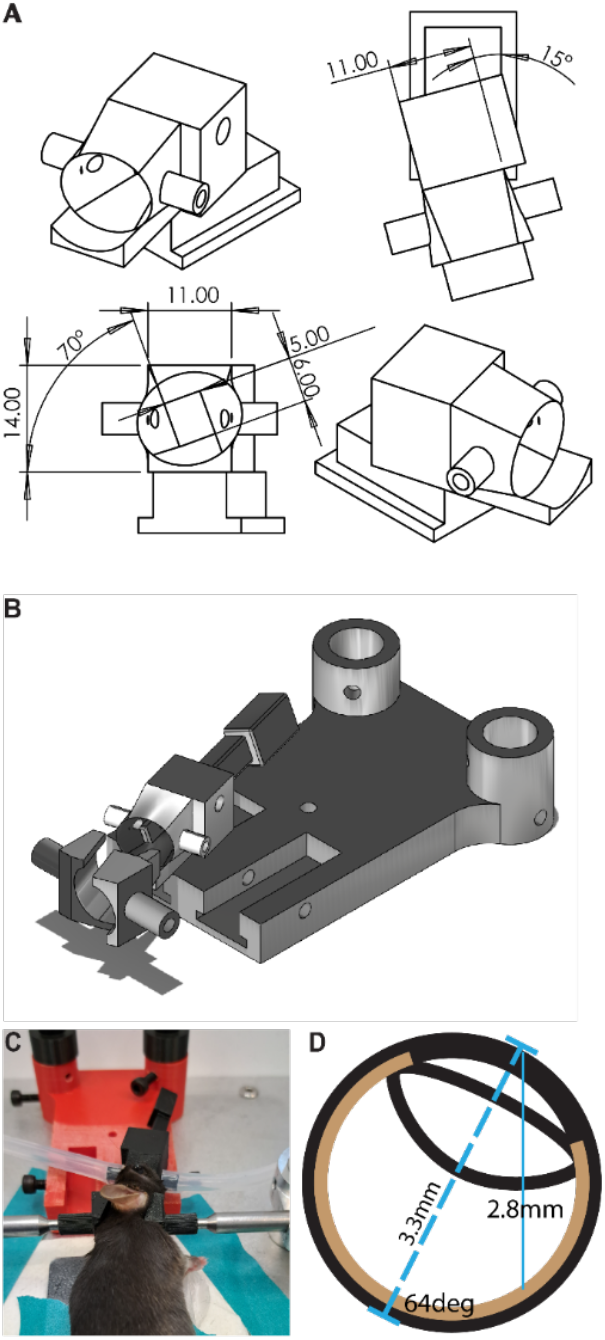
Custom Setup for Insonation of the Mouse Eye. **(A)** We designed a custom nosecone where the bitebar is rolled 70° and pitched 15° **(B)** The nosecone sits in a groove in a baseplate to allow for interchangeability of configurations so the right or left eye can face upward. Covers are included for the ear bars of the stereotactic stage used in the RK50. **(C)** When placed in the nosecone, the mouse’s eye faces up, the neck is fixed and supported. **(D)** A schematic of the eye when positioned in the nosecone: The net rotation aligns the visual axis with the sagittal plane with a 64° rotation outward. In this configuration, the retina is 2.8 mm inferior to the cornea.

### BRBO Volume and Success Increase with Pressure

Our first experiments applied FUS at a range of pressures to identify parameters which promoted BRBO without concurrent tissue damage at 4.667 MHz. We defined successful BRBO as EBD extravasation into retinal tissues. Experiments have determined that the success and safety of BBBO is dependent on mechanical index (MI)^15^, defined as the peak rarefactional pressure. (P_rp_) over the square root of the center frequency (f_c_) of the focused ultrasound: 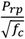, and thus we decided to test FUS-BRBO at various MI levels to find optimized parameters for opening and delivery. Representative fluorescence images of insonated and contralateral control eyes at each pressure show no opening at the lowest pressures in 15 mice (MI = 0, 0.2). At intermediate pressures (MI = 0.4, 0.6) we observed BRBO in 12 out of 24 mice. Finally at the two highest pressures (MI = 0.8, 1.0), we observed the opening in 11 out of 18 insonated eyes (**Fig. 2A**). We never observed BRBO in the contralateral (uninsonated) eye of any animal. Insonation at MI ranging from 0.2 to 1.0 showed a positive correlation of the FUS pressure with the surface area of EBD extravasation on the retinal flat mounts (R = 0.6172, p = 0.0002, linear regression, **Fig. 2B**). The success rate varied with the pressure and ranged between 43±13% at MI = 0.6 but increased to 100% at MI = 1.0 (p < 0.0001, two tailed unpaired heteroscedastic t-test, N = 14 and N = 4 respectively) (**Fig. 2C**).

**Figure 2.**
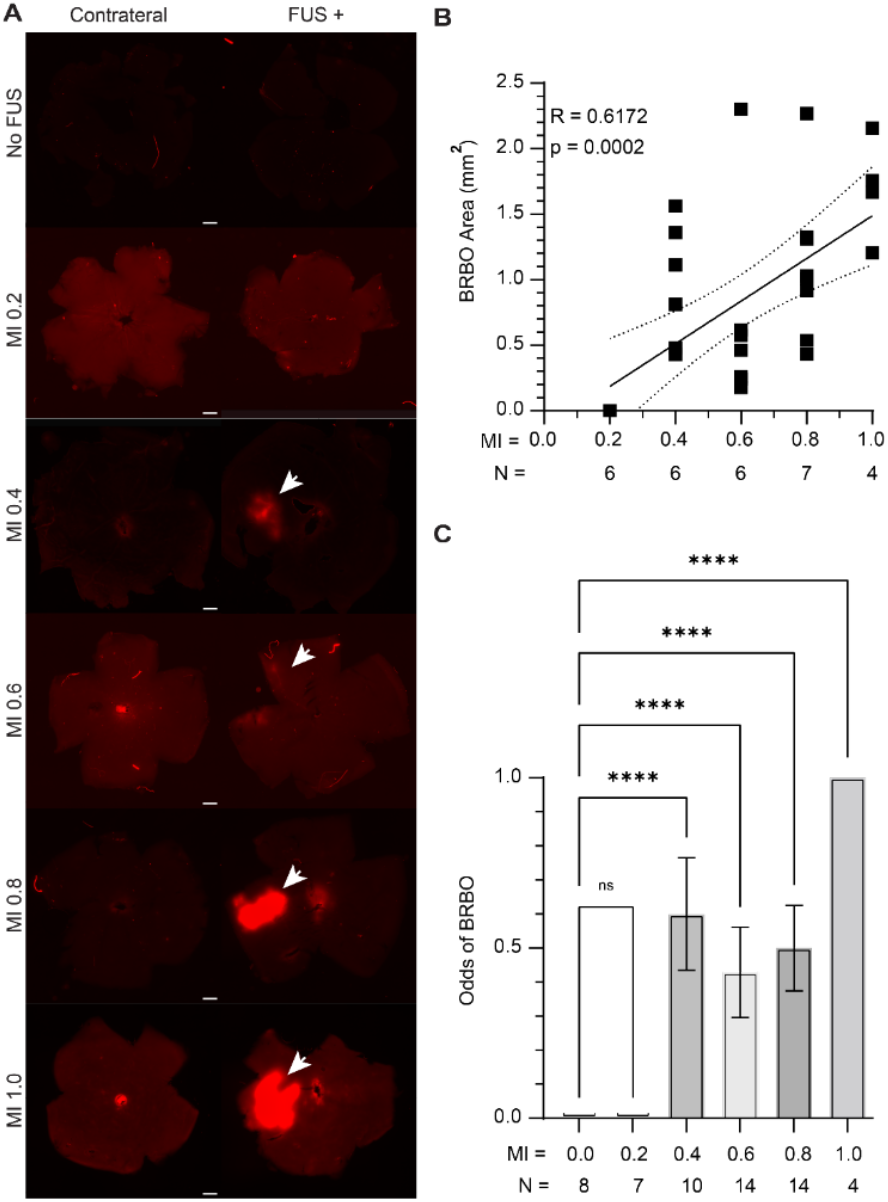
Magnitude and Odds of succesful BRBO are Dependent on FUS Pressure. **(A)** Representative images of BRBO from insonated eyes (FUS+) and controls (contralateral) at mechanical indices ranging from 0.0-1.0. EBD extravasation is visible in samples insonated at MI = 0.4 and above, with more intense EBD staining at the highest two pressures. Scale bar = 500 µm. **(B)** Increasing the pressure of the FUS causes a larger area of the BRB to be disrupted (R = 0.617, p = 0.0002, linear regression). Sample size for each group is listed below the data. Most cases of BRBO opened an area between 0.5 and 1.5 square millimeters. **(C)** Experimental success rates vary with pressure. At the highest and lowest pressures, there was complete success and failure, respectively. Error bars are standard deviation calculated as 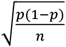, error bar is zero for MI = 0.0, MI = 0.2 and MI = 1.0. Sample size for each group is listed below the Figure (p < 0.0001, Ordinary one-way ANOVA with Tukey correction).

### Optimized Targeting Parameters Increases BRBO Success

For viral delivery with FUS-BRBO we intended to utilize the lowest possible pressure for BRBO but desired a higher success rate than 60±17% (**Fig. 3C**). Our comparison of BRBO at 4.667 MHz to 1.533 MHz insonation showed the lower frequency had higher odds of success at the same MI (p = 0.0001, unpaired two tailed heteroscedastic t-test, N = 10 and N = 5 respectively, **Fig. S1A**) which was not accompanied by an increase in BRBO area (p = 0.4268, unpaired two tailed homoscedastic t-test, N = 10 and N = 5 respectively, **Fig. S1B**). We hypothesized that this improved success was caused by the larger volume of FUS at a lower frequency leading to lower sensitivity to misalignment of ultrasound beam and the retina (**Fig. S1C**). To improve robustness of the targeting procedure, we optimized the protocol by changing the insonation from a single point on the retina (**Fig. 3A**) to multi-point spaced 200 µm apart in a vertical line along the axis of the transducer to effectively extend the area of effective insonation from the size of one full width half-maximum pressure area (0.3 x 1.7 mm) to 0.3 x 2.9 mm (**Fig. 3B**). This increased the success from 60%±17% among 10 mice when single targeting to 100% in a cohort of 7 mice when multi-point targeting (p < 0.0001, unpaired two tailed heteroscedastic t-test, **Fig. 3C**). This experiment was conducted at the lowest pressure where we saw BRBO, MI = 0.4. The representative opening is presented in **Fig. 3D**. This change in procedure did not cause an observed increase in the area of BRBO (p = 0.2526, unpaired two tailed homoscedastic t-test, **Fig 3E**).

**Figure 3.**
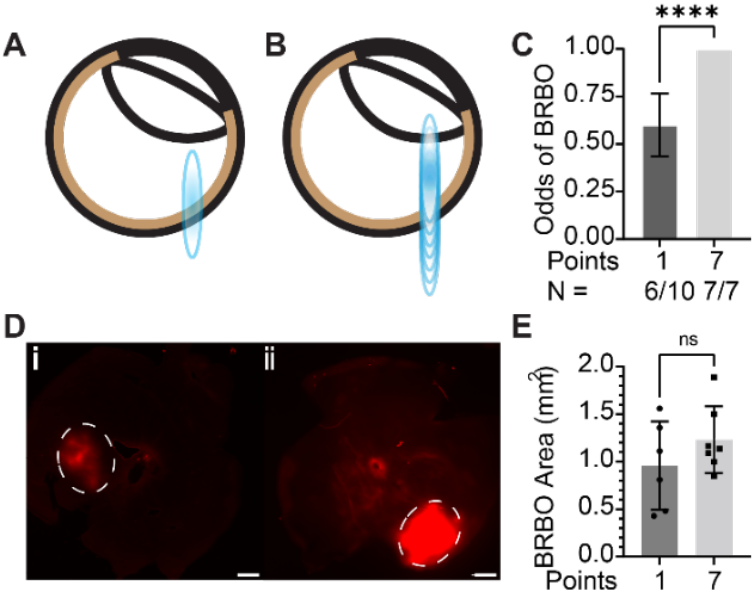
Multi-point Targeting Increases FUS-BRBO Success Rates. **(A)** The experiments described in Figure 2 targeted a single point below the cornea with FUS to cause BRBO. The size of the FUS beam overlaid on the retina is displayed here. **(B)** To increase the odds of success for FUS-BRBO we modified our targeting technique to incorporate multiple insonation points in a vertical line spaced by 200 µm. Visually, this is how the multi-point targeting was arranged around the retina. **(C)** All mice targeted with multipoint opening showed opening in the targeted eye, compared to the 6/10 targeted with a single point (p < 0.0001, unpaired two-tailed heteroscedastic t-test). **(D)** Representative images of the (i) single point and (ii) multipoint FUS-BRBO show opening within the circled area. Scale bar = 500 µm. **(E)** Opened area was not different when using single point or multi-point insonation when pressure was kept at MI = 0.4 (p = 0.2526, unpaired two-tailed homoscedastic t-test, N = 6 and N = 7 respectively).

### AAV.FUS.3 & AAV3 Preferentially Transduce Neurons in the Inner Retina Under FUS-BRBO

After verifying successful FUS-BRBO, we examined BRBO as a method to transduce retinal cells with two AAV serotypes – AAV.FUS.3 and AAV9. We selected MI = 0.4 for insonation as it was the lowest pressure with successful BRBO, aiming to reduce the potential for any side effects that can be seen in FUS-BRBO^39^ or FUS-BBBO at high pressure^15,19,40,41^. We then intravenously administered equivalent doses of AAV9 and AAV.FUS.3. The AAV.FUS.3 carried mCherry under CaG promoter, and AAV9 carried GFP under the same promoter to allow for comparison of their transduction under fluorescence microscopy (**Fig 4A-D**).

**Figure 4.**
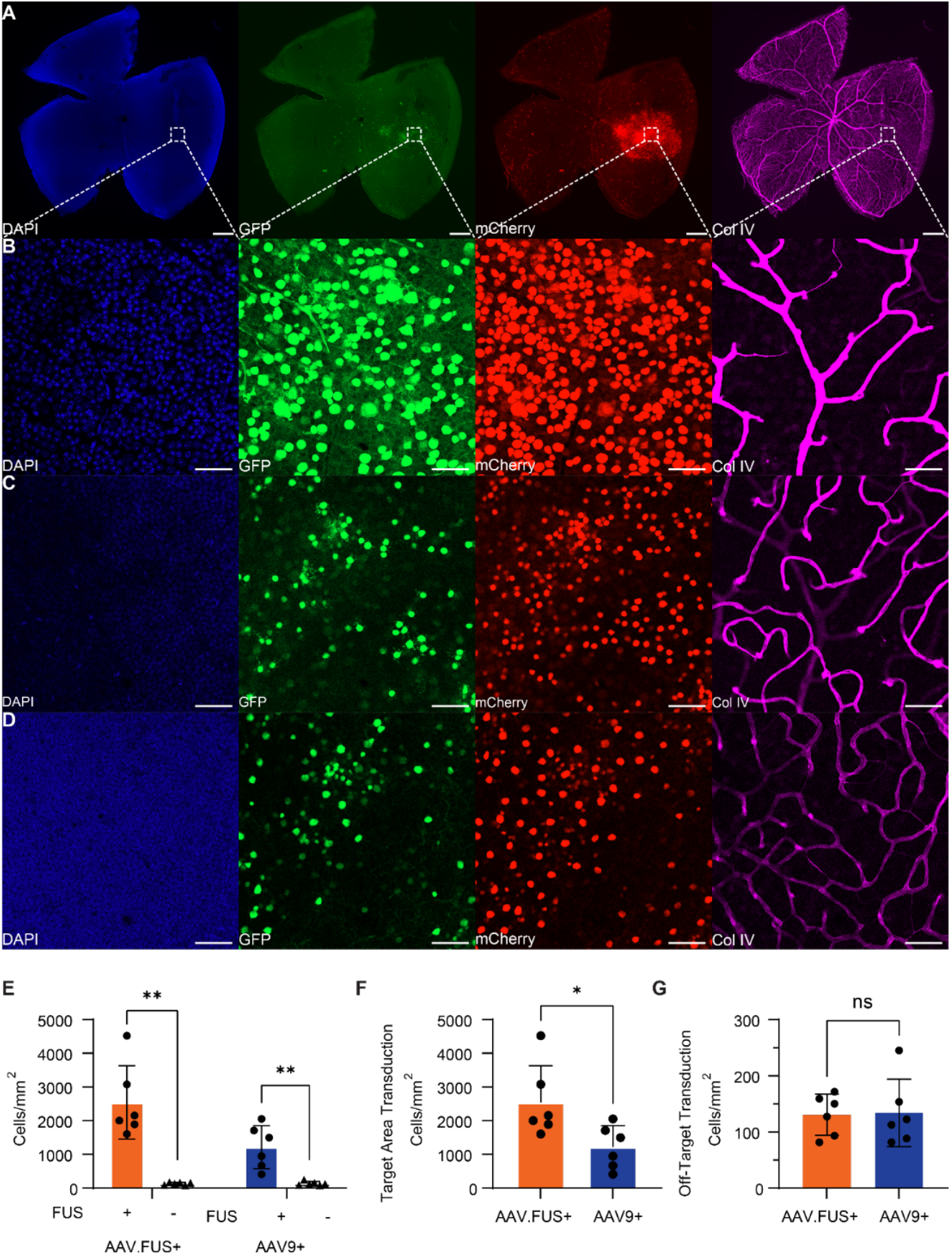
AAV.FUS.3 Transduces More Retinal Neurons than AAVG. **(A)** Flat mounted retinas show the area of insonation and increased transduction in the red and green channels (mCherry = AAV.FUS, GFP = AAV9). Scale bar = 500 µm. **(B-D)** Expanded insert shows the three vascular plexi and associated transduction of the inner retina. **(B)** Shows the superficial plexus, **(C)** shows the intermediate plexus, and **(D)** shows the deep plexus. **(E)** Application of FUS increased transduction when compared to untargeted regions of the same retina for both AAV.FUS.3 (19.5±2.7-fold; p = 0.0062, paired t-test, N = 6, error is SEM) and AAV9 (9.1±1.3 fold; p = 0.0068, paired t-test, N = 6, error is SEM). **(F)** Bulk transduction rates within the area of insonation show 2.1 times the number of cells expressing the AAV.FUS.3 carried fluorophore compared to AAV9 (p = 0.0115, paired t-test, N = 6). **(G)** This improvement is not present outside the area of insonation (p = 0.8590, paired t-test, N = 6).

When comparing the transduction in the insonated eyes to the contralateral control, AAV.FUS.3 transduction was improved 19.5±2.7-fold (p = 0.0028, two tailed unpaired homoscedastic t-test, N = 6, error is SEM) by FUS-BRBO, while AAV9 showed 9.1±1.3-fold (p = 0.0073, two tailed unpaired homoscedastic t-test, N = 6, error is SEM) (**Fig 4E**). The average transduction rate of AAV.FUS.3 in the insonated area on a flat mount was 2.1-fold higher than AAV9 (P = 0.0278, two tailed unpaired homoscedastic t-test, N=6, **Fig. 4F**). Specifically, AAV.FUS.3 transduced 2542±1091 cells per square millimeter of retina, while AAV9 transduced 1214±640 cells/mm^2^ (P = 0.0278, two tailed unpaired homoscedastic t-test, N = 6, error is standard deviation). When examining the area of the eye unexpos ed to FUS, both chosen vectors transduced comparable numbers of cells with AAV.FUS.3 transducing 131±37 cells/mm^2^ and AAV9 134±60 cells/mm^2^ (p = 0.9082, two tailed unpaired homoscedastic t-test, N = 6, error is standard deviation, **Fig. 4G**).

To examine the cell-type specificity of transduction with FUS-BRBO we next examined the transduction of non-neuronal cell-types in the retina—activated astrocytes (GFAP+), activated microglia (Iba1+), and Müller glia (Sox9+). Importantly, neither of the tested viral vectors transduced any of these non-neuronal cell types (**Fig. 5 A-C**).

**Figure 5.**
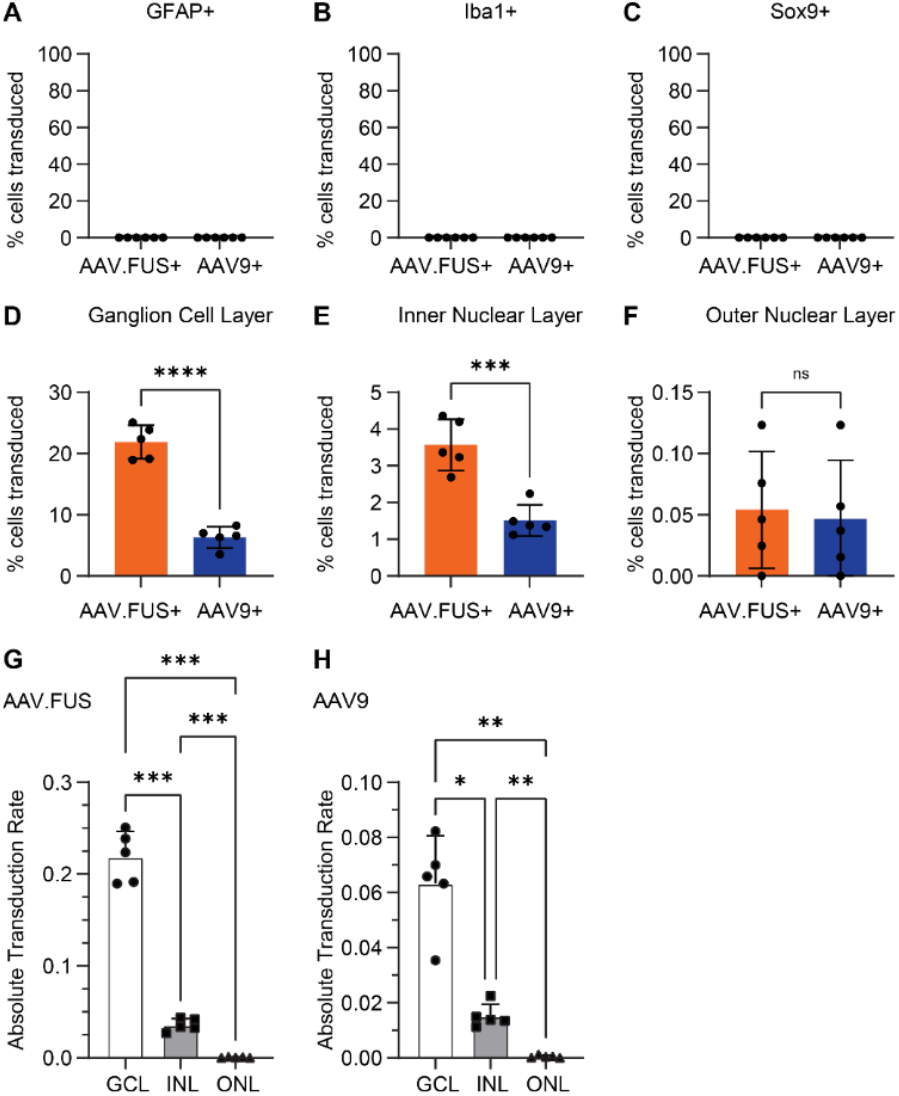
AAV. FUS.3 and AAVG Show Minimal Non-Neuronal Transduction and Primarily Transduce the Ganglion Cell Layer. **(A-C)**No transduction was seen in the three major groups of non-neuronal cells (activated astrocytes (GFAP+), microglia (Iba1+), or Muller glia (Sox9+)) (N = 6). **(D)** Transduction primarily occurred in the ganglion cell layer with 21.9±2.5% of nuclei costaining for AAV.FUS.3 and 6.3±1.5% of nuclei costaining for AAV9 (p < 0.0001). **(E)** Some transduction of cells in the inner nuclear layer (3.6±0.6% and 1.5±0.4% for AAV.FUS.3 and AAV9, respectively, p = 0.0005) and **(F)** negligible transduction of the outer nuclear layer (0.054±0.041% and 0.047±0.045% for AAV.FUS.3 and AAV9, respectively, p = 0.8106) was seen. **(G)** For AAV.FUS.3, GCL transduction was 6.3±0.3-fold higher than the INL (p = 0.0003, paired two tailed t-test, N = 5, error is SEM), 466±88-fold for GCL vs ONL (p = 0.0001, paired two tailed t-test, N = 4, error is SEM) (p < 0.0001, one way ANOVA, N = 5) and 80±15-fold for INL vs ONL (p = 0.0007, paired two tailed t-test, N = 4, error is SEM). **(H)** For AAV9 the changes were 4.5±0.4-fold for GCL vs INL (p = 0.0103, paired two tailed t-test, N = 5, error is SEM), 193±40-fold for GCL vs ONL (p = 0.0029, paired two tailed t-test, N = 4, error is SEM) (p = 0.0017, one way ANOVA, N = 5) and 53±16-fold for INL vs ONL (p = 0.0039, paired two tailed t-test, N = 4, error is SEM).

Next, we sought to explore tropism of AAV.FUS.3 and AAV9 in neuronal layers of the retina. In the ganglion cell layer (GCL), 21.9±2.5% of nuclei stained for AAV.FUS.3 and 6.3±1.5% of nuclei stained for AAV9 (**Fig. 5D**) (3.5±0.2-fold, p < 0.0001, paired two tailed t-test, N = 5, error is SEM). Transduction of the inner nuclear layer (INL) showed 3.6±0.6% for AAV.FUS.3 and 1.5±0.4% for AAV9 (2.4±0.1-fold, p = 0.0005, paired two tailed t-test, N = 5, error is SEM). Finally, the outer nuclear layer (ONL) showed a transduction rate of 0.054±0.041% for AAV.FUS.3 and 0.047±0.045% for AAV9 (p = 0.8106, paired two tailed t-test, N = 5, error is SEM). We then compared the transduction rates for each vector across the cell layers and found that the rate for the GCL was significantly higher than the other layers at 6.3±0.3-fold for GCL vs INL (p = 0.0003, paired two tailed t-test, N = 5, error is SEM) and 466±88-fold for GCL vs ONL (p = 0.0001, paired two tailed t-test, N = 4, error is SEM) for AAV.FUS.3 (p < 0.0001, one way ANOVA, N = 5). For AAV9, the changes were 4.5±0.4-fold for GCL vs INL (p = 0.0103, paired two tailed t-test, N = 5, error is SEM) and 193±40-fold for GCL vs ONL (p = 0.0029, paired two tailed t-test, N = 4, error is SEM) (p = 0.0017, one way ANOVA, N = 5).

### BRBO Did Not Produce Hemorrhages or Active Inflammation

Retinas that underwent FUS-BRBO and uninsonated contralateral controls were examined under brightfield microscopy to determine the presence of hemorrhage 20 minutes after the application of FUS. The 4X magnification brightfield imaging showed no visible hemorrhage in any eye at any pressure (104 eyes from 57 mice). A representative image of a retinal flat mount insonated at MI = 1.0 (**Fig. 6Ai**) shows no visible discoloration in the area affected by FUS (**Fig. 6Aii**). The contralateral eye is included for comparison (**Fig. 6B**).

**Figure 6.**
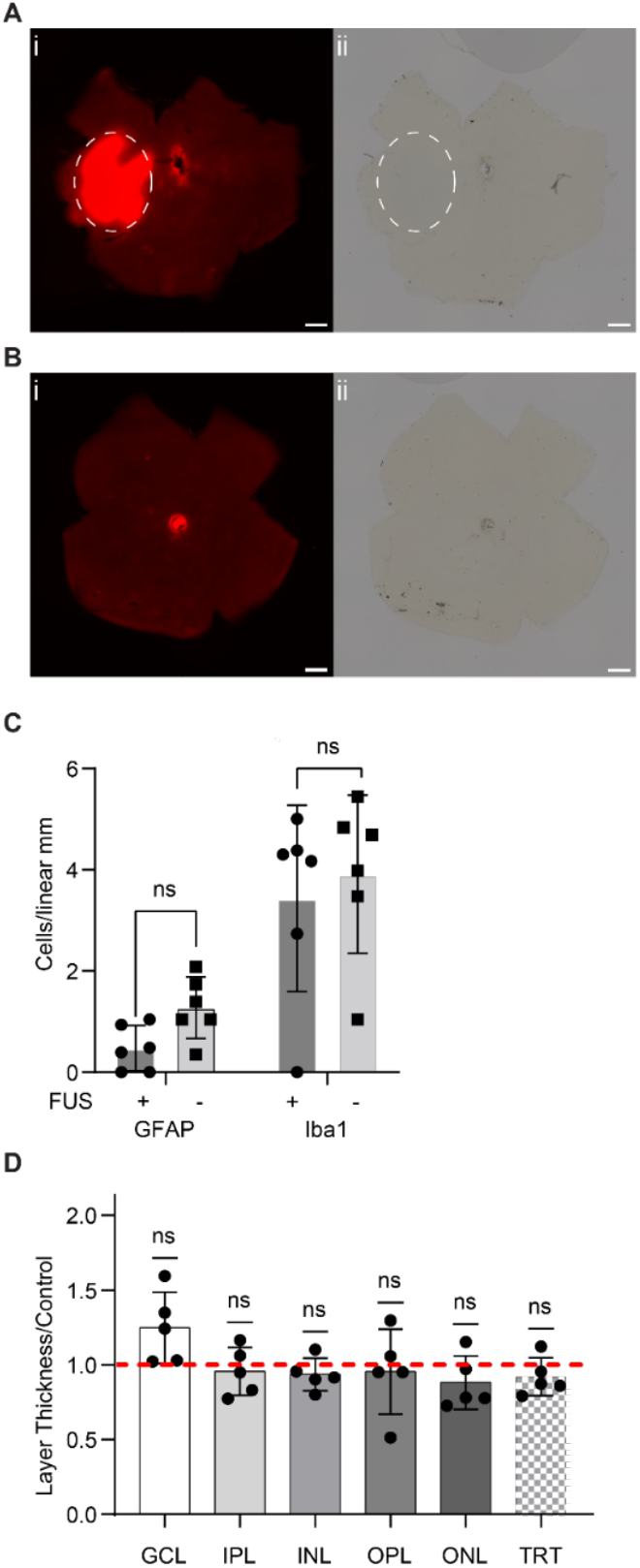
FUS-BRBO and Subsequent Viral Delivery are Safe. (**(A)**(i) At the highest pressure (MI = 1.0) in the area of opening (circle) retinas appear normal on brightfield imaging with no signs of hemorrhage or major disruption 20 minutes after FUS. (ii) Representative low power (4x) magnification image demonstrates this fact. Scale bar = 500 µm. **(B)** (i) Opening is not seen in the contralateral control eye, (ii) retinas also appear grossly normal. **(C)** Activated astrocytes (GFAP+) and microglia (Iba1+) were not found in the insonated area of the retina with greater frequency than in the same relative area of the contralateral eye (p = 0.0823, p = 0.03494, respectively, paired t-test, N = 6). **(D)** No significant difference was found between the thicknesses of the retina that was insonated and the same area of the contralateral control. (GCL = Ganglion Cell Layer p = 0.0817, IPL = Inner Plexiform Layer p = 0.5714, INL = Inner Nuclear Layer p = 0.2540, OPL = Outer Plexiform Layer p = 0.7333, ONL = Outer Nuclear Layer p = 0.2107, TRT = Total Retinal Thickness p = 0.2320, One sample t-test, N = 5).

The immunology and morphology of FUS-BRBO eyes was also investigated to verify the safety of this procedure at 3 weeks post insonation by staining. Retinal sections for astrocytic (GFAP+) and microglial (Iba1+) markers (**Fig 6C**). We found 0.5±0.4 GFAP+ cells per linear mm of section in the insonated area, and 1.3±0.6 GFAP+ cells per linear mm of section in the control eye, showing no significant elevation of GFAP due to insonation (p = 0.0823, paired t-test, N = 6) (**Fig. 6C**). Similarly, we found 3.4±1.8 Iba1+ cells per linear mm in the insonated area and 3.9±1.6 Iba1+ cells per linear mm in the uninsonated control retina, showing no significant difference between the groups (p = 0.3494, paired t-test, N = 6). To test for gross cell loss, we measured the thickness of the layers of the retina in the area of insonation and to a location matched control (contralateral) eye. No significant difference was seen in the thickness between the two eyes in any cell layer (GCL p = 0.0817, IPL p = 0.5714, INL p = 0.2540, OPL p = 0.7333, ONL p = 0.2107, TRT p = 0.2320, One-sample t-test, N = 5).

## DISCUSSION

In this study we designed a modified head stage to allow for stereotactic access to the eye (**Fig.1**). Using this stage we examined FUS-BRBO at 4.667 MHz for non-surgical delivery of a small molecule tracer, Evans Blue Dye (EBD), to the retina (**Fig. 2**). We achieved repeatable BRBO with FUS pressure at MI = 0.4 or higher. To maximize the potential of opening, we modified our FUS-BRBO procedure to incorporate multi-point targeting to extend the FUS beam (**Fig. 3**). We then investigated this technique’s ability to deliver viral vectors to the eye. We were successful in delivering both AAV.FUS.3 and AAV9 to the inner layers of the retina, seeing a 19.5±2.7-fold and 9.1±1.3-fold improvement in transduction from FUS, respectively (**Fig. 4**), which is consistent with prior results of this vector in the brain^42^. Both the insonation and transduction were not accompanied by inflammatory (**Fig. 5**) or structural (**Fig. 6**) changes to the retina, indicating this is a safe procedure.

Prior to this study FUS-BRBO had not been tested at frequencies greater than 1.1 MHz^37,39^. The retina is situated externally to the skull, reducing the need for avoiding signal attenuation from the bone that hampers targeting FUS to the brain^43^. Thus, retinal targeting is amenable to the use of higher ultrasound frequencies than those used in BBBO which reduces the diffraction-limited focusing and thus allows for greater spatial precision^44,45^. Interestingly, we did not see this effect of reduced opened area in our results (**Fig. S1B**). We did, however, observe differences in the chance of success at different frequencies (**Fig. S1A**), leading to our change in procedure prior to viral delivery experiments. The technique described in this study used external markers of the eye and stereotactic targeting to extrapolate the location of the retina, which will inherently be less precise than direct targeting methods such as MRI. Nevertheless, this technique could be combined with a direct imaging modality such as ultrasound to increase the accuracy of targeting subregions of the retina.

A key consideration for FUS-BRBO is the anatomical distinction between the inner and outer BRB. The inner BRB is formed by tight junctions between retinal vascular endothelial cells^8,46,47^. These vessels only supply the inner retinal layers. The outer BRB on the other hand is formed by retinal pigment epithelial cells and regulates exchange of macromolecules between the choroid and outer retina^6,7^. The results of this study indicate that FUS-induced BRBO predominantly affects the inner BRB, demonstrated here as 460 times the number of cells expressing AAV.FUS3-mCherry in the GCL as the ONL (**Fig. 4G,H**), suggesting that the degree of post-FUS permeability is not uniform across the retina^48^. Previous studies have shown that the vasculature of the retina are not uniformly susceptible to damage^49,50^, a similar predisposition may exist for FUS-BRBO, which may reflect some basic principle of the vascular biology of the retina. Importantly, this regional specificity may be advantageous for targeting diseases involving RGCs and inner retinal circuitry^51–53^. However, additional modifications will be needed to access the outer retina and photoreceptors^54,55^. Successful identification of the appropriate vectors and treatment-optimized ultrasound conditions would allow FUS-BRBO gene delivery to address a wider array of retinal pathologies^56–61^.

FUS-BRBO represents the potential for non-surgical alternative for delivery of gene therapies^62–65^ into the retina. Current standard of care for these patients is either intravitreal^66^ or subretinal^67^ delivery of viral vectors. These delivery modalities are limited by the inner limiting membrane^68^ or the need to produce a small retinal tear to access the subretinal space^69^ and the subsequent minor retinal detachment from subretinal fluid^70^. FUS-BRBO, conversely, has neither of these drawbacks, since vectors are administered systemically and no penetration is made into the eye. The potential adverse effects of systemic delivery of viral vectors can be mediated by designing specific vectors with enhanced specificity for target tissues^42^, and by using cell type-specific promoters to reduce peripheral expression^71^. We opted to use a generic promoter in this study to identify vector transduction of a variety of cell types within the retina^61^, still, we found primarily neuronal transduction (**Fig. 5**). Previous studies also evaluated systemic transduction of AAVs^72,73^ showing that primary target tissue *in vivo* is the liver, with AAV.FUS.3 showing nearly 7-fold reduction in transduction of liver compared to AAV9^42^.

Future studies should apply FUS-BRBO to disease models in animals to examine the safety and efficacy of FUS-mediated gene therapy delivery in specific retinal diseases. The non-surgical nature of this procedure is promising in clinical applications where ocular surgeries are expensive^74–76^, result in patient anxiety^12,75,76^, and are limited by physician availability^75–77^.

## MATERIALS AND METHODS

### Animals

C57Bl/6J (000664, Jackson Laboratory, Bar Harbor, ME) mice were purchased from Jackson Laboratory. Animals were housed according to a standard 12 h light/dark cycle with water and food ad libitum. Animal experiments were conducted under an approved protocol (no. IACUC-23-265-RU), in accordance with NIH guidelines, and followed the ARVO statement for the Use of Animals in Ophthalmic and Vision Science Research.

### Focused Ultrasound Blood-Retina Barrier Opening (FUS-BRBO)

Mice aged 9-14 weeks were anesthetized under 2.5% isoflurane in O_2,_ and intravenous access was gained via the lateral tail vein. Mice were moved to a modified stereotactic setup (**Fig. 1**) mounted in a Focused Ultrasound machine (RK50, FUS Instruments, Toronto, Canada) with a custom-designed 3D-printed mouse holder (**Supplementary Files 1-4**). The center of the cornea was identified with a targeting probe. A surgical drape was then placed over the mouse’s head with a hole over the eye. Sterile ultrasound gel was placed over the eye, a bath of degassed water was placed on top of the gel, and the FUS probe was placed in the degassed water.

Definity micro-bubbles (Perflutren Lipid Microsphere, Lantheus Medical Imaging, Billerica, MA, USA) were diluted in sterile saline (100um definity in 900um saline, Hospira, 00409-4888-10) to a final concentration of 2.9E12 microspheres/g and infused over the course of 10 seconds prior to insonation.

Ultrasound parameters were: 4.667 MHz center frequency with peak rarefactional pressures ranging from 0.425 MPa to 2.2 MPa. Pressures were based on calibration data provided by FUS Instruments (Toronto, Canada) and did not include any attenuation factors. A site 2.8 mm below the cornea (0,0,-2.8) was targeted using Morpheus software (FUS Instruments) and insonated 120 times with a 1 second period, 1% duty cycle.

### Evans Blue Dye Staining and Flat Mounts

After FUS-BRBO, animals were maintained in 2.5% isoflurane anesthesia and 50mg/kg of EBD (E2129-10g, Sigma-Aldrich, St. Louis, MO) was infused intravenously. Anesthesia was maintained for 20 minutes; mice were then sacrificed via 500mg/kg ketamine + 12.543 mg/kg xylazine intraperitoneal injection. Cardiac perfusion first with 10U/ml Heparin PBS (9041-08-1, Sigma-Aldrich), then 10% neutral buffered formalin (HT501128, Sigma-Aldrich) followed.

Both eyes were marked at the medial canthus with an electrocautery pen (927942, McKesson, Richmond, VA). Eyes were enucleated and placed in ice cold PBS. The cornea and lens were removed and 3 cuts were made in the eye cup. The sclera was removed from the retina with blunt forceps, and the retina was placed on nitrocellulose. Retinas were fixed in 10% formalin for one hour, removed from nitrocellulose, then washed in PBS three times for 30 minutes before storage at 4°C.

### BRBO Size Measurements

Fixed retinas were mounted on slides in Vectashield Antifade Mounting Medium with DAPI (H-1800, Vector Laboratories). Slides were then imaged on a BZ-X810 fluorescence microscope (Keyence, Osaka, Japan) at 4x magnification under brightfield and TxRed (560/40 nm excitation with 630/75 nm emission filtration) channels.

Fluorescence images of each retina were processed with the Meta Segment Anything Model (Meta AI, Menlo Park, CA). The location of BRBO was added to the model which then circumscribed the area of opening. The segmented region was processed in Matlab (MathWorks, Natick, MA) to compute the area.

### Transretinal Ultrasound Viral Delivery

Transretinal ultrasound viral delivery used the same protocol as FUS-BRBO with minor modifications. Before the microbubbles were administered, the mice were given a viral cocktail consisting of 10^10^ GC/g AAV.FUS.3 and 10^10^ GC/g AAV9. Viral payloads were nuclear localization sequence-tagged mCherry or GFP expressed under a constitutive CaG promoter (CAG-NLS-mCherry and CAG-NLS-GFP) to allow for differentiation between transduced cells. Flat mount experiments used AAV.FUS.3-mCherry and AAV9-GFP, while sectioning experiments used AAV.FUS.3-GFP and AAV9-mCherry.

### Flat Mount Histology

Mice were euthanized and their retinas extracted as detailed above. Once extracted and fixed, the retinas were then stained using the following protocol:

Retinas were blocked 10% goat serum (5425S, Cell Signaling Technology, Danvers, MA) in PBS++ (1xPBS (20-031-CV, Corning, Manassas, VA), 0.1% triton-x (X100-100ML, Sigma, Burlington, MA) and 0.1% Sodium Azide (S2002-100G, Sigma, Burlington, MA)) overnight at 4°C. Primary antibody staining consisted of 1:500 rat anti-mCherry (M11217, Invitrogen, Waltham, MA), 1:500 chicken anti-GFP (ab13970, Abcam, Waltham, MA), and 1:500 rabbit anti-Collagen IV (AB756P, Sigma-Aldrich) in 3% goat serum in 1xPBS, and allowed to incubate for three days on a shaker at 4°C. Retinas were washed for thirty minutes in PBS, three times, before secondary staining with 1:1000 DAPI, 1:500 goat anti-chicken AlexaFluor 488 (A-11039, Invitrogen), 1:500 goat anti-rat AlexaFluor 594 (A-11007, Invitrogen), 1:500 goat anti-rabbit AlexaFluor 647 (A-21245, Invitrogen) in 3% goat serum in PBS. Secondary stain was incubated overnight at 4°C on a shaker plate. Retinas were washed in PBS three times for thirty minutes before mounting in Antifade Mounting Medium (H-1800, Vector Laboratories).

Images were taken on the Keyence BZ-X810 fluorescence microscope (Keyence, Osaka, Japan) at 4x magnification for the purpose of identifying the areas of insonation. 20X magnification images were taken on a Zeiss LSM 800 confocal microscope using z-stacking. Five images were taken of the insonated area: one in the center, and four more at half the distance between the center and the edge of insonation.

### Cryosectioning and Staining

Mice underwent cardiac perfusion as before and eyes were marked at the medial canthus. Eyes were fixed for 4 minutes, the cornea and lens removed, and cryoprotected in 20% followed by 30% sucrose overnight. Eyecups were placed in OCT Compound (23-730-571, Fisher Healthcare, Houston, TX) in a 7mm x 7mm x 5mm mold with the burn marks facing upwards. The molds were directionally frozen by being placed on a metal block half submerged in liquid nitrogen.

Eyecups were serial sectioned across 15 slides on a cryostat (CM1950, Leica Biosystems, Nussloch, Germany). Sections were made at a 15 µm thickness in the axial plane.

Sections were stained on the slide using the following procedure: OCT was washed away with four five-minute washes in PBS++. Sections were blocked for one hour at room temperature 10% goat serum in PBS++, then incubated at room temperature in a primary antibody solution containing Anti-mCherry and Anti-GFP described above and one of either 1:500 rabbit anti-Iba1 (ab178846, Abcam), 1:500 rabbit anti-sox9 (#AB5535, Millipore Canada Inc, Etobicoke, Ontario, Canada), or 1:500 rabbit anti-GFAP Alexafluor 647 Conjugated (sc-33673, Santa Cruz Biotechnology, Dallas, TX). Sections were washed with PBS five times for five minutes each, secondary stain was applied, as before, the anti-rabbit antibody was removed when staining with anti-GFAP. Slides were again washed for five minutes five times before mounting in in Antifade Mounting Medium.

### Cell Counting and Histological Analysis

Section slides were imaged on the Keyance BZ-X810 and Zeiss LSM800 confocal for retinal layer thickness analysis and cell counting, respectively. Images of an equivalent area were taken on both retinas on slices taken an equivalent distance from the start of sectioning. ImageJ (NIH, Bethesda, MD) was used to calculate the thickness of the retina in the area of insonation as well as in the opposite eye in an equivalent location.

For cell counting, images were taken on an LSM800 of the same slides and locations and mCherry+ and GFP+ cells were counted in each cell layer in ImageJ. Cell densities in each cell layer were calculated by dividing the number of cells counted by the linear distance of the counted layer visible in the image. Positive cells in the far-red channel were also noted by layer, and the density was calculated in the same manner.

### Statistical Analyses

Statistical analysis was performed using Prism (GraphPad Software, version 9.2.0 (320)). A two-tailed *t*-test was used to compare two data sets, while one-way ANOVA with corrections for multiple comparisons was applied for comparisons involving more than two data sets. Single ratio data sets were compared against a hypothetical value of 1 with a two-tailed one sample T-test. Measurements are displayed with ± standard deviation unless specified. Statistical significance was set at \**P* < 0.05; \*\**P* < 0.01; \*\*\**P* < 0.001; and \*\*\*\**P* < 0.0001.

## ACKNOWLEDGEMENTS

Supported by National Eye Institute, National Institutes of Health, R21 Research Grant (R21EY032596), P30 Core Grant (P30EY002520), Individual Predoctoral Dual Degree Fellowship (F30EY037188); and an Unrestricted Award from Research to Prevent Blindness

