## Supplementary information for "Ultrasound-enhanced retinal delivery of engineered viral vectors"

### Supplementary Figures

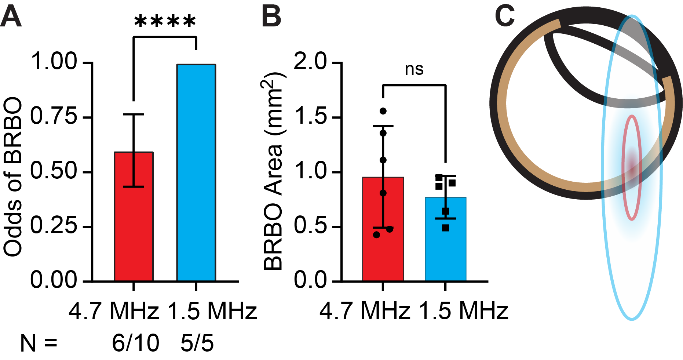

#### **Figure 1. 4.667 MHz and 1.553 MHz FUS Result Differential Success Rates, but Similar Opening Size**

(A) Using a reduced frequency of FUS (1.533MHz) increased the odds of success for BRBO (p = 0.0001, unpaired two tailed heteroscedastic t-test, N = 10 and N = 5 respectively) (B) but not the area of opening (p = 0.4268, unpaired two tailed homoscedastic t-test, N = 10 and N = 5 respectively). The only parameter varied in this experiment was the FUS frequency. Mechanical Index was held constant at 0.4, microbubble concentrations were the same, and number of FUS pulses and duty cycle were equivocal between experiments. (C) An overlay of the FUS beams on the eye shows the relative sizes of the 4.667MHZ (red) and 1.533 MHz (Blue).
